# Epigenetic analyses of planarian stem cells demonstrate conservation of bivalent histone modifications in animal stem cells

**DOI:** 10.1101/122135

**Authors:** Anish Dattani, Damian Kao, Yuliana Mihaylova, Prasad Abnave, Samantha Hughes, Alvina Lai, Sounak Sahu, Aziz Aboobaker

## Abstract

Planarian flatworms have an indefinite capacity to regenerate missing or damaged body parts owing to a population of pluripotent adult stems cells called neoblasts (NBs). Currently, little is known about the importance of the epigenetic status of NBs and how histone modifications regulate homeostasis and cellular differentiation. We have developed an improved and optimized ChIP-seq protocol for NBs in *Schmidtea mediterranea* and have generated genome-wide profiles for the active marks H3K4me3 and H3K36me3, and suppressive marks H3K4me1 and H3K27me3. The genome-wide profiles of these marks were found to correlate well with NB gene expression profiles. We found that genes with little transcriptional activity in the NB compartment but which switch on in post-mitotic progeny during differentiation are bivalent, being marked by both H3K4me3 and H3K27me3 at promoter regions. In further support of this hypothesis bivalent genes also have a high level of paused RNA Polymerase II at the promoter-proximal region. Overall, this study confirms that epigenetic control is important for the maintenance of a NB transcriptional program and makes a case for bivalent promoters as a conserved feature of animal stem cells and not a vertebrate specific innovation. By establishing a robust ChIP-seq protocol and analysis methodology, we further promote planarians as a promising model system to investigate histone modification mediated regulation of stem cell function and differentiation.

## Introduction

The promoters of developmental genes in mammalian embryonic stem cells (ESCs) are frequently marked with both the silencing H3K27me3 mark and active H3K4me3 marks. It has been proposed that this ‘bivalent’ state precedes resolution into full transcriptional activation or repression depending on ultimate cell type commitment (Bernstein et al. 2006; Voigt et al. 2013; Harikumar and Meshorer 2015). The advantage is that bivalency represents a poised or transcription-ready state, whereby a developmental gene is silenced in ESCs, but can be readily rendered active during differentiation to a defined lineage. Evidence for this comes from the finding that 51% of bivalent promoters in ESCs are bound by paused polymerase (RNAPII-Ser5P), compared with 8% of non-bivalent promoters (Lesch and Page 2014; Brookes et al. 2012); demonstrating a strong but not complete association. Bivalency may also protect promoters against less reversible suppressive mechanisms, such as DNA methylation (Lesch and Page 2014). Bivalent chromatin has also been discovered in male and female germ cells at many of the gene promoters that regulate somatic development, and may underpin the gametes’ ability to generate a zygote capable of producing all cellular lineages (Yamaguchi et al. 2013; Cui et al. 2009; Hattori et al. 2013; Sachs et al. 2013; Lesch et al. 2013; Lesch and Page 2014).

It remains unclear whether the poised bivalent promoters of developmental genes are an epigenetic signature of vertebrates or arose earlier in the ancestor of all animals. Recently, the orthologs of bivalent genes that sit at the top of transcriptional hierarchies in mammalian development, were also found to be poised in chicken male germ cells (Lesch et al. 2016). Sequential ChIP has also established H3K4me3/H3K27me3 co-occupancy of promoters in zebrafish blastomeres (Vastenhouw et al. 2010). Conversely, comparatively few bivalent domains were identified in Xenopus embryos undergoing the midblastula transition (Akkers et al. 2009). Xenopus genes which appear to have signals for both H3K4me3 and H3K27me3 originate from cells in distinct areas of the embryo, and as such the observed bivalency can be explained by cellular heterogeneity (Akkers et al. 2009).

Given that bivalency correlates with pluripotency in ESCs, planarian pluripotent adult stem cells or neoblasts (NBs) represent one possible scenario where poised promoters could have an important role in invertebrates, if this regulatory feature is conserved. Planarian NBs are a population of adult dividing cells that collectively produce all differentiated cells during homeostatic turnover and regeneration (Aboobaker 2011; Rink 2013). Several RNA-binding proteins, such as *piwi* and *vasa*, typically associated with nuage of germ cells are also expressed in planarian NBs where they function in the maintenance of pluripotency (Reddien et al. 2005; Palakodeti et al. 2008; Solana 2013; Shibata et al. 2016; Lai and Aboobaker 2018). Moreover, the ability of NBs to differentiate upon demand must also require well-regulated transcriptional and epigenetic processes, and poised, bivalent promoters may constitute an effective way of coordinating the differentiation of these stem cells.

Here we develop an optimized ChIP-seq methodology for planarian NBs and combine this with informatics approaches to establish robust approaches for studying histone modifications at transcriptional starts sites (TSSs). This enabled us to identify genes with inactive/low expression in the NB population, but with greatly increased expression in post-mitotic NB progeny that are actively differentiating, with bivalent promoters in planarian NBs by combining transcriptomic and epigenetic analyses. Our findings indicate that bivalent promoters in pluripotent stem cells are not just a facet of vertebrates, but may have a role in regulating pluripotency in embryonic and adult stem cells across animals.

## Results

### Genome-wide annotation of transcribed loci in asexual *Schmidtea* mediterranea genome and categorization by proportional expression in FACS populations

We sought to produce an annotation of all transcribed loci on the asexual *Schmidtea mediterranea* genome (SmedAsxl v1.1) utilising both *de novo* assembled transcriptomes and 164 independent RNA-seq datasets covering RNAi knockdown-, regenerating-, whole worm-, and cell compartment-specific datasets (**Figure 1A, Supplementary File 1**). The inclusion of these diverse datasets was to improve the overall representation of the genome, and is useful for discovering potential non-coding RNAs and protein-coding genes expressed at low levels, both of which may not have been fully covered by individual studies limited by read number, or reliant on homology based annotation processes such as MAKER (Cantarel et al. 2008).

**Figure 1:**
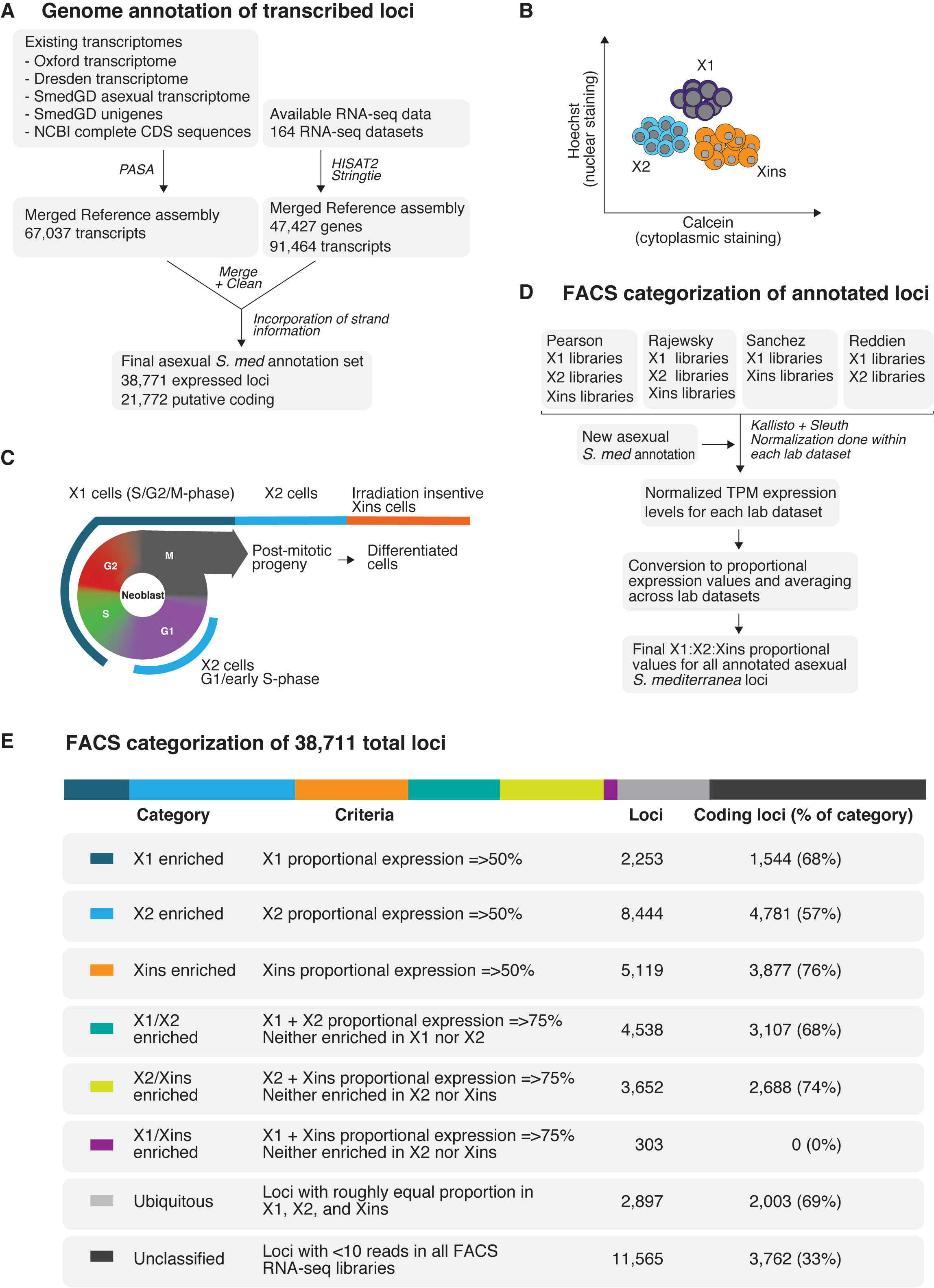
**A**. Overview of methodology for annotating the *Schmidtea mediterranea* asexual genome based on expression. 164 RNA-seq datasets, 4 *de novo* transcriptome assemblies, NCBI complete CDS sequences, and Smed Unigenes were mapped to the SmedGD Asxl v1.1 genome. Reference assemblies were merged, cleaned to remove potential splice variant redundancies, and the best representative transcript for each genomic locus was chosen. Strand information was obtained by BLAST to Uniprot, prediction of longest ORF and data from strand-specific libraries. This process yielded a total set of 38,771 loci. **B**. Methodology for gating X1, X2, Xins cell populations based on nuclear to cytoplasmic ratio during Fluorescent Activated Cell Sorting (FACS). **C**. Diagram depicting how X1, X2, Xins FACS population gates relate to cell cycle and differentiation stage. **D.** Overview of methodology for categorization of annotated loci based on FACS RNA-seq datasets. FACS RNA-seq datasets were mapped to our annotated genome using Kallisto and normalized with Sleuth. Normalization was done individually for each of the lab’s datasets. Normalized TPMs were converted to proportions between available FACS categories of each lab, and a final consensus X1:X2:Xins proportion was calculated. **E.** A table presenting number of loci in different FACS classification groups, as well as number of protein-coding genes in each group based on Transdecoder evidence.

Our new expression-based annotation identified 38,711 expressed loci, 21,772 of which are predicted to be coding (**Figure 1A**). Moreover, compared to the current available annotation of the *Schmidtea mediterranea* asexual genome (Smed GD 2.0) (Robb et al. 2008), our annotation discovered 10,210 new potential protein coding loci that are expressed at similar overall levels to previously annotated protein coding genes. A total of 6,300 genes from the existing MAKER homology based annotation were not present in our expression driven annotation. Further analysis of these MAKER-specific genes shows that they generally have no or very little potential expression within the 164 RNA-seq libraries utilised for our annotation (**Supplementary Figure 1**).

In the absence of transgenic approaches and antibodies for confirmed cell lineage markers, Fluorescence Activated Cell Sorting (FACS) gating cell populations stained with Hoechst and calcein is the best available tool for isolating NBs, progeny, and differentiated cells (Hayashi et al. 2006; Romero et al. 2012). FACS allows for two irradiation sensitive compartments to be discerned: the ‘X1’ gate representative of S/G2/M-phase NBs with >2C DNA content; and the ‘X2’ gate representative of G1 phase NBs and post-mitotic progeny with 2C DNA content. The third FACS population, ‘Xins’, represents an irradiation-insensitive population with a higher cytoplasmic to nuclear ratio (**Figure 1B and 1C**). These cell compartments are heterogeneous, with subpopulations of NBs expressing epidermal, gut and other lineage-specific markers present within the X1 population (Scimone et al. 2014; Wurtzel et al. 2015; Van Wolfswinkel et al. 2014), and the X2 compartment consisting of an amalgam of G1 NBs and lineage-committed post-mitotic progeny (Baguñá and Romero 1981; Hayashi et al. 2006; Zhu et al. 2015; Molinaro and Pearson 2016).

We used the publicly available RNA-seq datasets for these three different FACS populations in order to compare the expression of our annotated loci in these three distinct compartments (Önal et al. 2012; Labbé et al. 2012; Van Wolfswinkel et al. 2014; Zhu et al. 2015; Duncan et al. 2015). We first looked at the normalized TPM expression levels for annotated loci in our annotated genome in the FACS population datasets originating from four different planarian labs **(Supplementary Figure 2).** This revealed a rough congruence between different FACS populations from different labs (**Supplementary Figure 3A**).

We transformed absolute TPM expression values into proportional values for each FACS compartment in each of the datasets (**Figure 1D; Supplementary Figure 3B**). These proportional values were then averaged across datasets, to produce a final set of X1:X2:Xins proportions for 27,206 loci (18,010 of which are predicted to be protein-coding) that had at least 10 reads mapped in at least one FACS RNA-seq library.

We were now able to sort all annotated genes according to whether their predominant expression (i.e. => 50% expression) is in X1 (S/G2/M-phase NBs), X2 (NBs and stem cell progeny) or Xins (differentiated cells) (**Figure 1E**). We confirmed this analysis by Gene Ontology (GO) analyses and verification of the proportional expression profiles for known genes previously described as being enriched in X1, X2 or Xins (**Supplementary Figure 4**) (Solana et al. 2012; Önal et al. 2012; Labbé et al. 2012). We also re-analysed FACS single-cell RNA-seq datasets in the context of our genome annotation, and visualisation of the data by breakdown into our defined FACS expression categories was entirely consistent with this data (**Supplementary Figure 5**). In particular we note that those genes with the highest proportion of expression the X2 compartment are indicative of genes expressed in post-mitotic undifferentiated NB progeny, with only very little expression in NBs themselves **(Supplementary Figure 5)**.

Together these analyses provide a set of annotations and expression values that are directly related to the genome assembly, allowing integration of ChIP-seq data to investigate correlations between epigenetic marks and gene expression in the different planarians cell FACS compartments.

### An optimized ChIP-seq protocol reveals H3K4me3 and H3K36me3 levels correlate with gene expression in planarian NBs

Research into the epigenetic mechanisms governing stem cell pluripotency in planarian NBs is still in its infancy (Dattani et al. 2018). Previous work has uncovered a lack of endogenous DNA methylation in the *Schmidtea mediterranea* genome, and characterized loss of function phenotypes for members of the NURD complex (Scimone et al. 2010; Jaber-Hijazi et al. 2013; Vásquez-Doorman and Petersen 2016), COMPASS and COMPASS-like families (Hubert et al. 2013; Duncan et al. 2015; Mihaylova et al. 2017) The first study to utilize ChIP-seq in planarians documented the effects of *mll1/2* and *set1* RNAi with respect to the activation mark H3K4me3 (Duncan et al. 2015). However, we revisited this data and noted that the total number of ChIP-seq reads from -1 million X1 sorted NBs was comparatively low in comparison to those from *Drosophila melanogaster* S2 ‘carrier’ cells.

We developed an optimized ChIP-seq protocol for FACS sorted X1 NBs without the addition of excess ‘carrier’ cells. Instead, a -3% *Drosophila* ‘S2’ spike-in was added to our chromatin before immunoprecipitations (IP) simply as a method to normalize any technical differences in IPs across our replicate libraries (Orlando et al. 2014). We were able to generate high quality uniquely mapped reads to our annotated *Schmidtea mediterranea* genome using only 150-200,000 X1 cells per IP – 5 to 7 times less material than the previously established planarian protocol (Duncan et al. 2015). With our protocol, *Drosophila* ‘spike-in’ reads accounted for an average of 27% of X1 H3K4me3 libraries compared to an average of 87% in the previous study’s X1 H3K4me3 libraries. Moreover, *Drosophila* ‘spike-in’ reads accounted for 9% of our X1 H3K36me3 libraries compared with 99% of the single X1 H3K36me3 replicate included in a previous study (Duncan et al. 2015) (**Supplementary Figure 6**).

We tested the robustness of our ChIP-seq protocol with reference to both H3K4me3 and H3K36me3 – epigenetic marks that are known to positively correlate with gene expression in other model systems. H3K4me3 is laid down by the trithorax group (trxG) complexes containing SET or MLL enzymes at active promoters near TSSs (Bledau et al. 2014; Denissov et al. 2014; Sirén et al. 2014; Hu et al. 2013). H3K36me3 is a mark of transcriptional elongation, and is deposited on histones as they are displaced by RNA polymerase II and as such this modification is enriched towards the 3’ end of genes (Wagner and Carpenter 2012; Li et al. 2002). H3K36me3 is hypothesized to prevent spurious transcriptional initiation at cryptic promoter-like sequences within exons and, in yeast, this is achieved by the recruitment of histone deacteylase complexes (HDAC) that erases elongation-associated acetylation (Carrozza et al. 2005; Joshi and Struhl 2005).

As predicted, ChIP-seq of H3K4me3 in X1 NBs revealed a high average peak around the TSSs of genes characterized as being X1 enriched (**Figure 2A**). Conversely, we observed comparatively lower H3K4me3 deposition at the TSSs of Xins enriched genes not expressed or expressed only at very low levels in X1 cells. Intermediate levels of H3K4me3 in the X2 compartment are consistent with this FACS population being a mixture of NBs and post-mitotic progeny. Indeed, genes with the highest proportion of X2 expression (i.e. ‘high ranking X2 genes’) indicative of expression in post-mitotic progeny but not NBs had lower levels of H3K4me3 in X1 cells compared with low ranking X2 genes that retain expression in cycling NBs **(Figure 2B).** A base by base Spearman’s Rank correlation of ChIP-seq signal to FACS proportional expression values of annotated loci across a 2.5 kb region either side of the TSS, shows a positive correlation between genes defined by high X1 proportional expression and the levels of H3K4me3 deposition close to the TSS (**Supplementary Figure 7A**). On the other hand, there is a negative correlation between H3K4me3 deposition and genes with high Xins proportional expression across the same region. Thus, a high H3K4me3 ChIP-seq signal reflects higher expression of a locus in X1 NBs, whereas lower H3K4me3 signal reflects lower X1 NB gene expression but higher expression in the differentiated Xins compartment.

**Figure 2:**
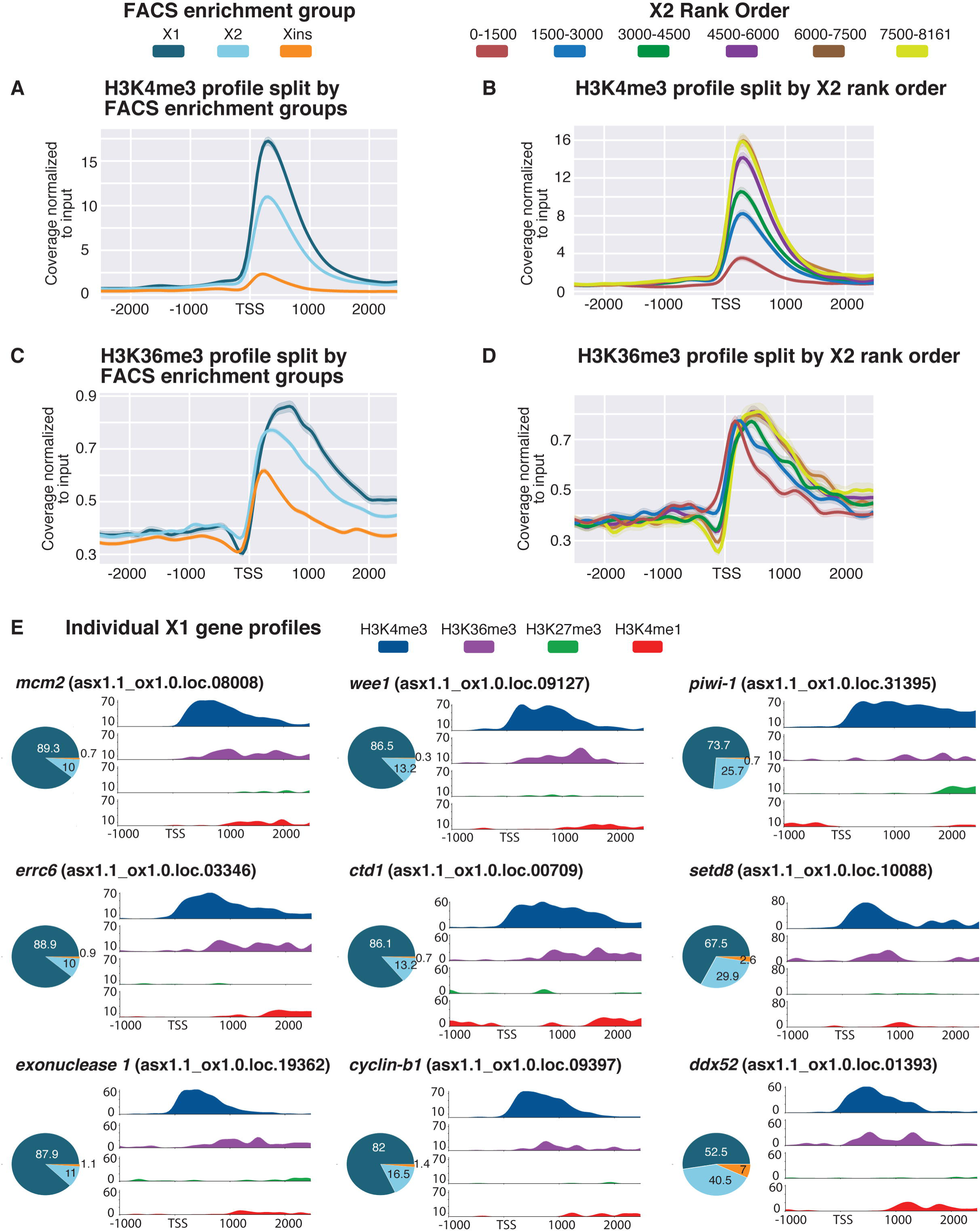
Histone marks for actively transcribed genes in X1 NBs. **A**. Average H3K4me3 ChIP-seq coverage profiles across X1, X2 and Xins enriched loci in X1 NBs across biological replicates following outlier removal. Y-axis represents the difference in coverage between sample and input. X-axis represents 2.5kb up- and downstream of the TSS. H3K4me3 signal is highest around promoter-proximal region close to the TSS for X1 enriched loci in NBs consistent with the role of H3K4me3 in active transcription. **B.** H3K4me3 ChIP-seq profiles following outlier removal for X2 genes ranked from high to low X2 proportional expression. H3K4me3 signal in NBs decreases with an increase in proportion of X2 expression, indicative of high-ranking X2 genes having a predominant role in post-mitotic progeny as opposed to NBs. **C.** Average H3K36me3 ChIP-seq profile across X1, X2 and Xins enriched loci in X1 NBs across biological replicates following outlier removal. Y-axis represents the difference in coverage between sample and input. X-axis represents 2.5kb upstream and downstream of the TSS. Signal for H3K36me3 is promoter-proximal for Xins genes, whereas the magnitude of signal is greater and shifted 3’ for X1 genes. **D.** H3K36me3 ChIP-seq profiles following outlier removal for X2 genes from high to low X2 proportional ranking. H3K36me3 signal in NBs shifts to the 3’ end with a decrease in X2 proportion, consistent with these lowly-ranked genes having transcriptional activity in NBs. **E.** H3K4me3 and H3K36me3 (active marks) and H3K4me1 and H327me3 (suppressive marks) ChIP-seq profiles for highly-expressed X1 genes in NBs. Y-axis represents percentage coverage for each mark and allows for the 4 epigenetic marks to be directly compared. X-axis represents 1.0 kb upstream and 2.5 kb downstream of the TSS. Pie Charts represent proportional expression for each gene in X1 (dark blue), X2 (light blue) and Xins (orange).

ChIP-seq plots of H3K36me3 split by FACS gene expression revealed, as predicted, a higher average peak around X1 enriched genes when compared with the X2 and Xins FACS enrichment categories (**Figure 2C**). Importantly, the average peak for X1 genes is located towards the 3’ end of genes, whereas the smaller Xins peak is promoter-proximal by comparison. This can be explained by a higher level of transcriptional elongation of X1 transcripts in NBs compared with Xins genes that have a predominant expression in the differentiated compartment. When splitting X2 enriched genes by rank order, we observe that genes with highest expression in the X2 compartment and, as a consequence lowest transcript abundance in NBs, have an enrichment for H3K36me3 at the promoter-proximal end of the gene (**Figure 2D**). Conversely, with decreasing X2 proportional expression and a concomitant increase in transcriptional activity in the NB compartment, the average peak of H3K36me3 is shifted downstream of the TSS towards the 3’ ends of genes.

We also looked at the individual H3K4me3 and H3K36me3 profiles of genes known to be highly expressed in NBs, and compared this to the signal for the suppressive marks H3K4me1 and H3K27me3 (see later). We confirmed that known metazoan genes associated with stem cell maintenance, such as cell-cycle and replication related genes (i.e. *mcm2, cyclin-B1, wee1, ctd1*), RNA-binding proteins (*piwi-1, ddx52*), DNA-damage response (DDR) genes (*errc6-like, exonuclease 1*) and epigenetic-related genes (*setd8-1*), all have high levels of H3K4me3 at the promoter-proximal end and H3K36me3 in the gene body, but a comparatively low signal for the suppressive marks H3K4me1 and H3K27me3 (**Figure 2E**).

### Levels of repressive histone marks H3K27me3 and H3K4me1 at TSSs in NBs correlate with gene expression

Utilising our optimized ChIP-seq protocol, we investigated the occurrence of two additional histone modifications: H3K27me3, a repressive promoter mark catalysed by the PRC2 complex, and H3K4me1, a mark mediated by the MLL3/4 family of histone methyltransferases that correlates both with active enhancers and inactive promoter regions (Cheng et al. 2014; Calo and Wysocka 2013).

Genes that are categorized as being X1 enriched have low levels of H3K27me3 deposition at the TSS, compared with Xins enriched genes which are silenced in NBs (**Figure 3A**). A positive correlation is observed between the level of H3K27me3 and expression in the Xins compartment in a window from the TSS to 1kb downstream. This fairly broad domain of H3K27me3 deposition is consistent with previous studies in mammals (**Supplementary Figure 7b**) (Hawkins et al. 2011; Pauler et al. 2009). Conversely, a negative correlation at the TSS is observed between H3K27me3 signal and genes with high X1 expression (**Supplementary Figure 7b**). Consequently, the genome wide pattern for H327me3 is the opposite to that observed for H3K4me3. When splitting X2 genes by rank we note that genes with higher transcriptional enrichment in the post-mitotic compartment have a higher overall level of H3K27me3 at the promoter proximal region compared to genes that have NB expression (**Figure 3B)**.

**Figure 3:**
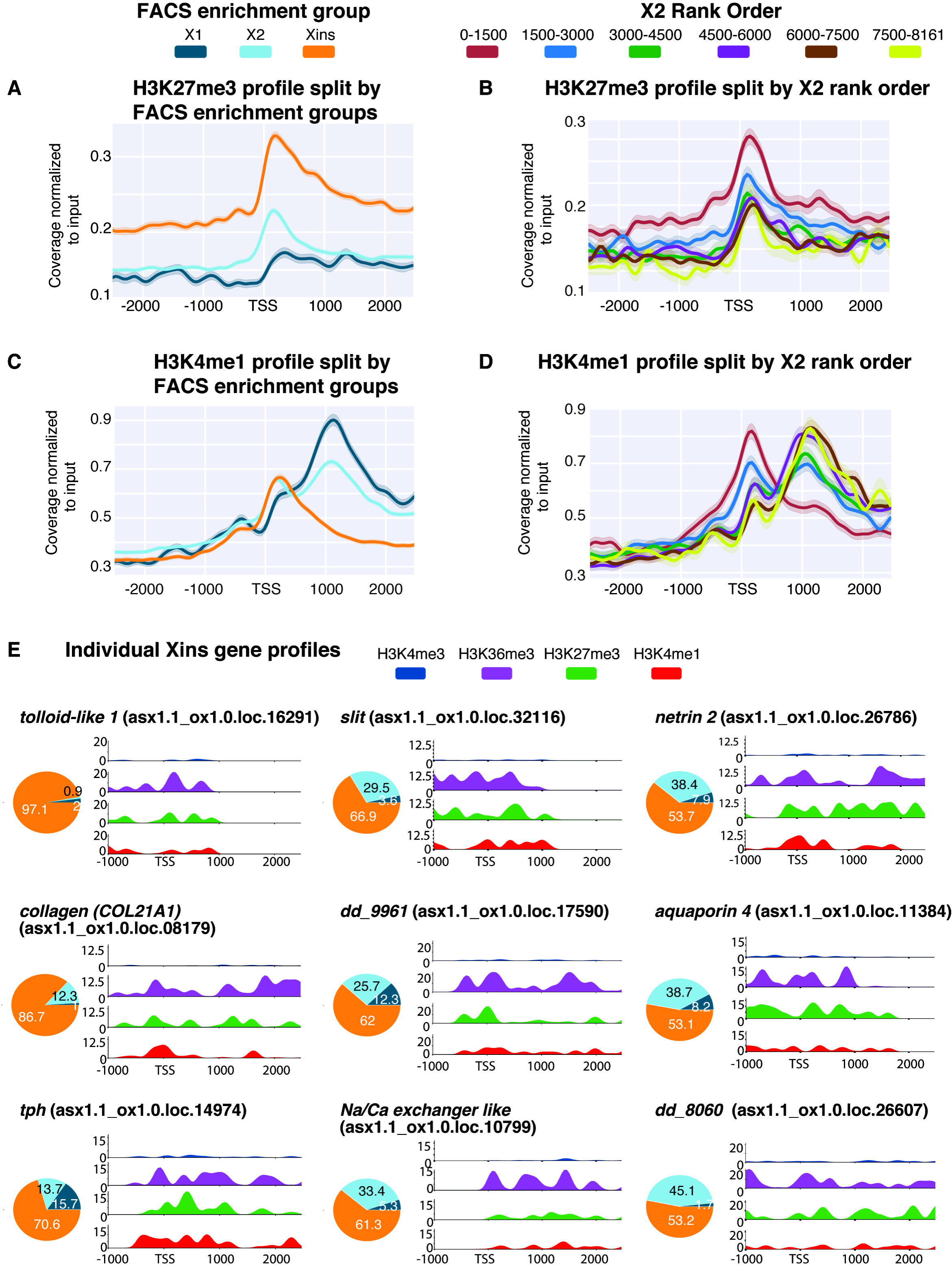
Histone marks for inactive genes in X1 NBs. **A**. Average H3K27me3 ChIP- seq profile across X1, X2 and Xins enriched loci in X1 NBs across 3 biological replicates following outlier removal. Y-axis represents the difference in coverage between sample and input. X-axis represents signal 2.5kb upstream and downstream of the TSS. **B.** H3K27me3 ChIP-seq profiles following outlier removal for X2 genes from high to low X2 proportional ranking. H3K27me3 signal increases with an increase in proportion of X2 gene expression, indicative of these high-ranking X2 genes being transcriptionally silenced or lowly expressed in NBs. **C.** Average H3K4me1 ChIP-seq profiles following outlier removal across X1, X2 and Xins enriched loci in X1 NBs. Y-axis represents the absolute difference in coverage between sample and input. X-axis represents signal 2.5kb upstream and downstream of the TSS. **D.** H3K4me1 ChIP-seq profiles following outlier removal for X2 genes from high to low X2 proportional ranking. Highly ranked X2 genes have a H3K4me1 signal at the promoter-proximal region, and a decrease in X2 ranking coincides with a peak shift-1kb downstream of the TSS. **E.** H3K4me3, H3K36me3, H3K4me1 and H327me3 NB ChIP-seq profiles for highly-expressed Xins genes. The Y-axis scale represents percentage coverage for each mark. X-axis represents 1.0kb upstream and 2.5 kb downstream of the TSS. Pie Charts represent proportional expression for each gene in X1 (dark blue), X2 (light blue) and Xins (orange).

The distribution of the H3K4me1 mark is noticeably different compared to that observed for either H3K27me3 or H3K4me3. Specifically, Xins loci have high levels of H3K4me1 at the TSS in X1 NBs, consistent with these genes being expressed at low levels in NBs, whereas X1 loci have H3K4me1 peaks that are on average-1kb downstream of the TSS **(Figure 3C)**. This data suggests that the H3K4me1 signal shifts away from the TSS for genes that are actively expressed in NBs, in agreement with previous observations in mammals (Cheng et al, 2014). Further evidence of this peak shifting comes from analysis of X2 enriched genes sorted by rank order of expression (**Figure 3D**). Highly ranked X2 genes are marked with H3K4me1 at the promoter-proximal region. As the proportion of X2 enrichment decreases, indicative of increasing expression the G1 NB compartment, the average H3K4me1 profile becomes bimodal, eventually shifting downstream of the TSS (**Figure 3D**).

We plotted the epigenetic profiles of individual genes known to have high Xins proportional expressions and that have validated expression patterns both by single-cell RNA sequencing data and *in situ* hybridisations (**Figure 3E**) (Fincher et al. 2018; Plass et al. 2018). For example, these genes are expressed almost exclusively in the muscle (*COL21A1, slit1*), parenchyma (*glipr1, tolloid-like 1*), cathepsin+ cells (*dd961, aquaporin 1*), non-cilliated neurons (*tph*, *dd8060*), and protonephridia (*Na/Ca exchanger-like*), all have high H3K27me3 signal at the TSS consistent with these genes being silenced in NBs. Moreover, these Xins enriched genes all have a high H3K4me1 signal at the TSS that anti-correlates with H3K4me3 deposition, in support of an earlier hypothesis that H3K4me1 limits the role of H3K4me3 interacting proteins (Cheng et al. 2014). We also observe an atypical placement of H3K36me3 at the TSS of individual Xins genes which supports the previous suggestion that that H3K36me3 may silence genic loci when placed at a promoter-proximal region of a gene (Wu et al. 2011).

### Correlations of H3K7me3 and H3K4me3 profiles against FACS proportions provide evidence for promoter bivalency in NBs

Having demonstrated that known active and suppressive marks correlate with gene expression in planarian NBs, we investigated whether promoter bivalency could act to keep genes in a poised state prior to the onset of differentiation. Bivalent promoters are characterized by the presence of both the activating mark H3K4me3 and repressive mark H3K27me3. The simultaneous presence of both these marks keeps the gene in a poised transcriptional state, with low or no expression, and upon differentiation resolves such that only one of the two marks is dominant. We reasoned that loci that are off or have relatively low proportional expression in X1 NBs, but which are upregulated during the differentiation process in post-mitotic progeny (high X2 expression), would be good candidates for potential regulation by bivalent promoters in NBs. Additionally, in the absence of sequential or co-ChIP-seq technologies for planarians, using genes no or very low expression in NBs greatly reduces the likelihood that any bivalent signals are due to cell heterogeneity. This is because these genes would not be expected to have high levels of H3K4me3 in any (or at least very few cells) in the X1 NB compartment.

We plotted the percentage of maximum coverage for both H3K4me3 and H3K27me3 for the top 1000 genes for each of the three FACS enrichment categories (**Figure 4A-C**). A plot for the top 1000 X1 genes shows that these genes have a higher level of H3K4me3 compared to H3K27me3 (**Figure 4A**), whereas the top 1000 Xins genes have on average a much higher H3K27me3 signal compared to H3K4me3 (**Figure 4C)**. Consistent with our hypothesis, the top 1000 X2 genes, have peaks that are of similar magnitude for both of these functionally opposing epigenetic marks (**Figure 4B)**.

**Figure 4:**
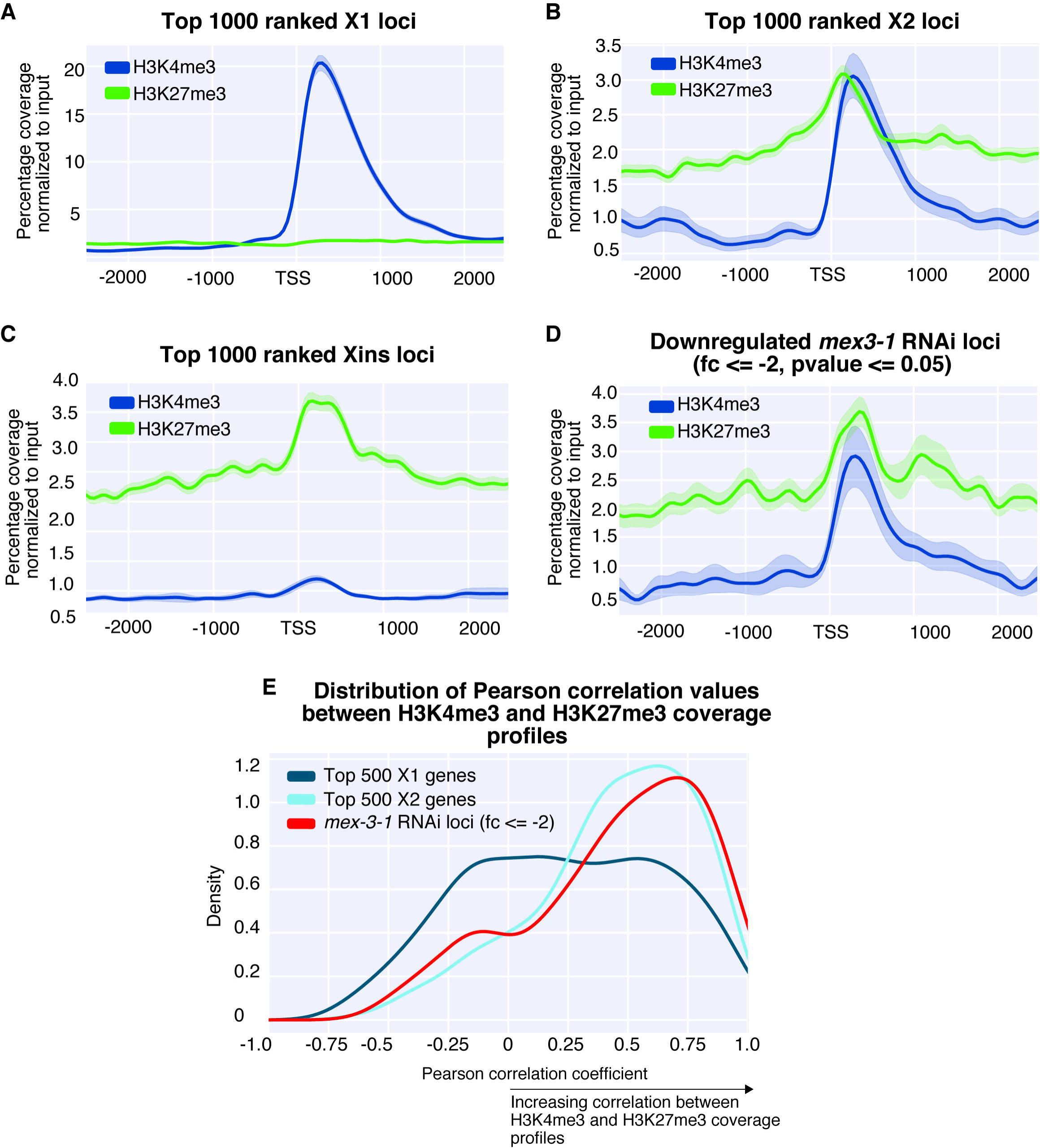
A-E: Average H3K4me3 and H3K27me3 ChIP-seq profiles in X1 NBs across 3 biological replicates. Y-axis is percentage coverage after normalization to input to allow both ChIP-seq profiles to be directly compared. Plots are shown for: **A.** Top 1000 ranked X1 genes by expression. **B.** Top 1000 ranked X2 genes. **C.** Top 1000 ranked Xins genes. **D.** 285 *Smed-mex3-1* down-regulated loci with >2-fold change (p<0.05). **E.** A distribution of Pearson correlation values for top 500 X1, X2, and 285 =>2-fold downregulated *mex-3-1* downregulated loci. The Pearson Correlation coefficient was calculated between the H3K4me3 and H3K27me3 values at each 50bp window starting from -1000bp downstream of the TSS and 1500bp upstream of the TSS.

We also plotted the epigenetic profiles of genes that are downregulated following RNAi of the planarian homolog of the RNA-binding protein MEX3 (Zhu et al. 2015). Previously, *mex3-1* has been shown to be necessary for generating the differentiated cells of multiple lineages, and consistent with a role in the differentiation process we found that the downregulated genes (downregulated 2-fold; p-value <=0.05) had a higher average X2 proportional expression value (62.4%) compared with that of X1 (12.5%) (**Supplementary File 2**). As expected, we note a paired H3K4me3 and H3K27me3 ChIP-seq signal for these *mex3-1* downregulated genes (**Figure 4D)**.

One possibility is that that our observations are a result of genes with high X2 expression only having the H3K4me3 mark only whilst other genes exist in a H3K27me3-only state in NBs. This would produce an average profile that appears bivalent when many genes are looked at simultaneously. To rule out this possibility, we plotted the distribution of Pearson correlation coefficients between H3K4me3 and H3K27me3 for the top 500 ranked X1, X2 and 285 *mex3-1(RNAi)* downregulated loci. This showed a strong positive correlation between H3K4me3 and H3K27me3 for top 500 X2 loci and *mex3-1* downregulated loci, compared to a weak or no average correlation for X1 loci (**Figure 4E**). This is consistent with the interpretation that bivalency is present at promoters of genes that are highly enriched for expression in the X2 compartment.

### Planarian orthologs to mammalian bivalent genes are marked by H3K4me3, H3K27me3 and paused RNA Pol II at the promoter-proximal region

RNA Polymerase II (RNAPII) pausing at genes that are highly inducible has been hypothesized to play a pivotal role in preparing genes for rapid induction in response to environmental or developmental stimuli. In a number of mammalian cellular contexts, bivalent genes have been shown to have a high density of paused RNA Pol II at the promoter-proximal region compared to genes which are actively transcribed, therefore allowing genes to be maintained in a transcriptionally poised state (Stock et al. 2007; Ferrai et al. 2017; Liu et al. 2017). Paused RNA Pol II can be distinguished from other forms by a phosphorylation at Ser5 (Ser5P) of the YSPTSPS heptad repeat at the C-terminus of the largest subunit of the Pol II complex. This heptad repeat is conserved across metazoans (Corden 2013), and is found in *S. mediterranea*.

ChIP-seq for RNAPII-Ser5P in NBs revealed that X2 enriched genes have a higher level of paused RNA Pol II at the promoter proximal region compared to X1 genes (**Figure 5A**). More significantly, highly ranked X2 genes with high expression in post-mitotic progeny and little expression in NBs have the highest amount of paused RNA Poll II close to the TSS, and with increasing expression in NBs the enrichment for this mark decreases (**Figure 5B**).

**Figure 5:**
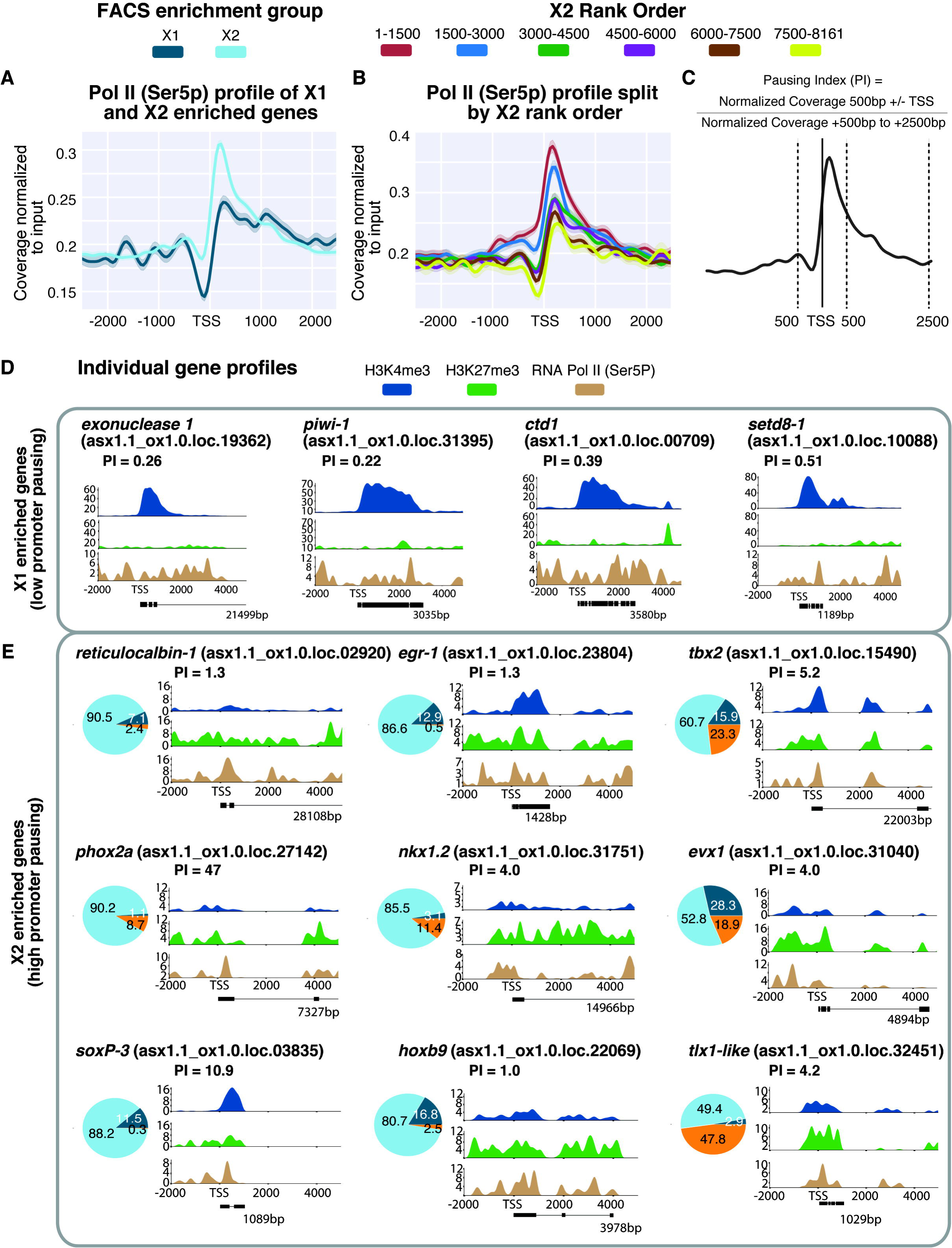
**A.** Average paused RNAPII-Ser5P ChIP-seq profile across X1 and X2 enriched loci in X1 NBs across biological replicates following outlier removal. Y-axis represents the difference in coverage between sample and input. X-axis represents signal 2.5 upstream and downstream of the TSS. **B.** RNAPII-Ser5P ChIP-seq profiles following outlier removal for X2 genes from high to low X2 proportional ranking. RNAPII-Ser5P signal increases with an increase in proportion of X2 gene expression, indicative of these high-ranking X2 genes being transcriptionally silenced but maintained in a permissive state for rapid induction. **C.** Calculation for Pausing Index (PI) of genes => 1kb. We divided normalized coverage between +/- 500bp TSS by normalized coverage +500bp to +2.5 kb. For genes under 2.5 kb, we inspected RNAPolI-Ser5P profiles visually to confirm whether Pol II pausing was enriched at the promoter-proximal region. **D.** Individual profiles for H3K34me3 and H3K27me3 of highly enriched NB X1 genes. X1 genes have a high level of H3K4me3 and levels of H3K27me3 correspond to intron regions and are not enriched at the promoter-proximal region. RNAPII-Ser5P signal is not enriched at the promoter-proximal region compared with the gene body, and as a result PI <1. **E.** We selected highly enriched X2 genes with a PI => 1 that have both H3K4me3 and H3K27me3 enriched at the promoter-proximal region, together with an enrichment of RNAPII-Ser5P close to the TSS. Pie Charts represent proportional expression for each gene in X1 (dark blue), X2 (light blue) and Xins (orange).

We calculated the pausing index (PI) for all annotated genes in our genome that have a total annotated length of ≥ 1kb. For our particular genome annotation, we calculated the PI as the read coverage (normalized to input) +/- 500bp either side of the annotated TSS divided by the normalized read coverage from +500bp to +2500bp from the TSS (**Figure 5C**). We applied a conservative definition of a gene as being significantly stalled for transcription if the PI ≥ 1. As expected, individual genes highly expressed in the NB compartment had both a low PI and were not enriched for RNAPII-Ser5P at the promoter-proximal region, thereby confirming our methodology was accurate at the gene level (**Figure 5D**). We also found that X2 genes with high PI scores had, on average, higher Pearson correlation coefficients between H3K4me3 and H3K27me3 (indicative of a bivalent state) compared with both X1 and X2 genes that have lower PI scores (**Supplementary Figure 8**). Given this correlation, we chose individual X2 enriched genes with high PI values and plotted the ChIP-seq profiles for H3K4me3, H3K27me3 and RNAPII-Ser5P as a percentage of maximum coverage for each mark.

Amongst genes enriched for these three signatures of bivalent promoters were those that have orthology to transcription factor (TF) families and include the Hox (*hoxb9*), Nkx (*nkx1.2*), Even-skipped (*evx-1*), Paired-like (*phox2A*) and T-Gbox (*tbx2*) and Tlx (*tlx1-like*) gene classes (**Figure 5E**). Indeed, previous studies in both mouse ESCs (Bernstein et al. 2006) and quiescent muscle stem cells (Liu et al. 2013) have shown that members of these gene families are typically marked by both H3K4me3 and H3K27me3. A paired level of these marks at the TSS for these individual genes suggests the existence of bivalent chromatin states at these conserved developmental genes and confirms our correlational analysis of X2 loci (**Figure 4E**). Moreover, single-cell sequencing data and pseudotime analyses plots made from single cell data show that these genes are expressed at detectable levels in very few, if any, *smedwi-1+* cells (the archetypal NB marker) and are instead enriched in post-mitotic cells of specific lineages (**Supplementary Figure 9**) (Fincher et al. 2018; Plass et al. 2018).

One caveat of our analyses is that the bivalent profiles of X2 enriched differentiation related genes may, for some individual genes that appear bivalent, reflect admixture of transcriptionally active and repressed states within the X1 NB compartment. For example, previous work has shown that the X1 compartment is highly heterogeneous with subsets of *piwi-1+* NBs expressing lineage specific TFs (Van Wolfswinkel et al. 2014). These genes, such as *SoxP-3* and *egr-1,* which are in fact X2 enriched according to our dataset and others (Labbé et al. 2012), appear to have a paired H3K4me3 and H3K27me3 signal (**Figure 5E**). Given that they are known to be expressed in a subset of cells in the X1 compartment and are definitive markers of lineage-primed NB subsets that will go through one more cell division (as validated by *in situ* hybridisation, condensin knockdown studies (Van Wolfswinkel et al. 2014; Lai et al. 2018) and single-cell RNA-seq data (Wurtzel et al. 2015; Plass et al. 2018; Fincher et al. 2018) no definitive conclusions concerning bivalency of these particular genes can be reached.

## Discussion

In this study, we have produced a *Schmidtea mediterranea* asexual genome annotation based on gene expression, and integrated FACS RNA-seq datasets from different laboratories to calculate consensus proportional expression values for each annotated locus in the X1, X2 and Xins cellular compartments. We have developed an optimized ChIP-seq protocol, and employed this to generate robust genome-wide profiles of the active H3K4me3 and H3K36me3 marks and repressive H3K4me1 and H3K27me3 marks in planarian NBs.

We find that the active marks H3K4me3 and H3K36me3, and suppressive H3K4me1 and H3K27me3 marks in X1 NBs correlate with the proportion of total transcript expression of these loci in X1 cells, validating our NB ChIP-seq methodology. These analyses showed that genes associated with stem cell differentiation, and which are expressed at low levels in X1 population but activated at high levels in the X2 population, are marked with both H3K4me3 and H3K27me3 marks at comparable levels at the TSS. Moreover, these genes were also highly marked with paused RNA polymerase (RNAPII-Ser5P) at the promoter region consistent with the definition of transcriptionally poised bivalent genes. Although we cannot entirely rule out cell heterogeneity within the X1 NB population as a factor contributing to our observation of promoter bivalency, our focus on both genes with high X2 expression (post-mitotic NB progeny) and orthology to vertebrate transcription factors known to have bivalent profiles, provide strong evidence that bivalent histone marks may be involved in poising of genes for activation upon NB commitment and differentiation in planarians.

The existence of promoter bivalency in invertebrates, prior to our work here, has been contentious. For example, the mammalian orthologs of bivalent genes in *Drosophila* germ cells were found to have only repressive H3K27me3 deposited at their promoters (Schuettengruber et al. 2009; Gan et al. 2010; Lesch et al. 2016). However, in a more recent study using fly embryos, the Pc-repressive complex 1 (PRC1) that binds to H3K27me3 was shown to co-purify with both the Fsh1 (ortholog of mammalian BRD4) that binds to acetylated histone marks and to Enok/Br140 (orthologs to subunits of mammalian MAZ/MORF histone acetyltransferase complex). ChIP-seq identified two groups of PRC1/Br140 genomic binding sites that were either defined by strong H3K27me3 signal or strong H3K27ac signal (i.e. actively transcribed genes). Both groups were also marked with narrow peaks of H3K4me3 at the TSS (Kang et al. 2017). These recent findings also argue for the existence of bivalent-like promoters outside of vertebrates, at least with respect to the binding of chromatin regulatory complexes, and extends the model to suggest that acetylation may be important in the resolution of bivalent protein complexes during development.

One key role of bivalency is thought to be to allow the maintenance of pluripotency in ESCs, by having genes involved in differentiation and commitment both silent but competent to switch on if the right signals are received. Our data suggest that this mechanism is likely to be important for pluripotency in planarian pluripotent NBs, as genes that can switch on rapidly upon differentiation appear to be the bivalent. Indeed, these genes also included planarian orthologs to mammalian TFs that have been documented to be bivalent in ESCs. Consequently, we are able to present a case for promoter bivalency in planarian NBs and in doing so demonstrate that this process is not necessarily vertebrate-specific. This novel finding adds to the growing body of evidence which suggests a deep conservation of regulatory mechanisms involved in stem cell function (Juliano et al. 2010; Alié et al. 2015; Solana 2013; Solana et al. 2016; Lai and Aboobaker 2018) as well as combinatorial patterns of post-translational modifications (Schwaiger et al. 2014; Sebé-Pedrós et al. 2016; Gaiti et al. 2017). Epigenetic studies in the unicellular relative of metazoans, *Capsaspora owczarzaki*, could not find any evidence of bivalency given the absence of H3K27me3, and epigenetic studies in the sponge *Amphimedon queenslandica* (Gaiti et al. 2017) and the cnidarian *Nematostella vectensis* (Schwaiger et al. 2014) have also not revealed any evidence for this approach to gene regulation. Further work will be required to establish when bivalent chromatin evolved in animals.

Overall our development of a robust ChIP-seq protocol for use with planarian sorted NBs, together with good coverage for four definitive and essential epigenetic marks, establishes a resource for the future planarian studies investigating the epigenetic regulation of stem cell function.

## Materials and Methods

### Reference assembly and annotations

Previous transcriptome assemblies (Oxford (ox_Smed_v2), Dresden (dd_smed_v4), SmedGD Asexual, Smed GD Unigenes) were downloaded from and PlanMine (Brandl et al. 2016) and Smed GD 2.0 (Robb et al. 2008). NCBI complete CDS sequences for *Schmidtea mediterranea* were also downloaded. Sequences were aligned to the SmedGD Asexual 1.1 genome with GMAP (Wu and Watanabe 2005) and consolidated with PASA. An independent reference assembly was also performed by mapping 164 available RNA-seq datasets with HISAT2 (Sirén et al. 2014) and assembly was performed with StringTie. The PASA consolidated and StringTie annotations were merged with StringTie.

An intron jaccard score (intersection of introns / union of introns) was calculated for all overlapping transcripts. Pair-wise jaccard similarity scores of 0.9 or greater were used to create a graph of similar annotations. From the resultant cliques of transcripts, one was chosen to be the representative transcript for the locus, by prioritizing transcript length, ORF length, and BLAST homology.

Strand information for annotations was assigned by utilising in house strand-specific RNA-seq libraries, BLAST homology, and longest ORF length. Transdecoder was run utilizing PFAM and UNIPROT evidence to identify protein-coding transcripts in the annotations. Detail methods are recorded in an IPython notebook (**Supplementary File 3**).

### FACS proportional expression value generation for annotated loci

Kallisto (Bray et al. 2016) was used to pseudo-align RNA-seq libraries originating from four labs (Önal et al. 2012; Labbé et al. 2012; Van Wolfswinkel et al. 2014; Zhu et al. 2015; Duncan et al. 2015) (**Supplementary Figure 2**) to our expression-based annotation of the asexual *S.mediterranea* genome. This generated TPM values for each annotated locus. Sleuth was used to calculate a normalization factor for each library. For each locus, the TPM values of member transcripts (potential isoforms) were summed to generate a consensus TPM value and then normalized accordingly. Replicates within each lab dataset were then averaged.

Normalized TPM values for each lab dataset were converted to a proportional value as a representation of expression in FACS categories. We next calculated three sets of pairwise ratios (X1:X2, X1:Xins, X2:Xins) using these proportional values. Given two of the three ratios, a third ratio can be ‘predicted’. Consequently, we calculated 3 ‘observed’ ratios and 3 ‘predicted ratios’. A good Spearman’s rank correlation was observed for the X2:Xins ratio and as such we kept these observed proportions, and calculated an inferred X1 proportion. Detailed methodology is documented in **Supplementary File 4** and full list of X1,X2, and Xins proportional values is available in **Supplementary File 5**.

### ChIP-seq protocol

For each experimental replicate, 600’000-700’000 planarian X1 cells were isolated, (sufficient for ChIP-seq of 3 histone marks and an input control) by utilisation of a published FACS protocol (Romero et al. 2012). We dissociated cells from an equal number of head, pharyngeal, or tail pieces from 3-day regenerating planarians. For whole worm ChIP-seq, wild-type worms were starved for 2 weeks prior to dissociation.

Following FACS, cells were pelleted. The pellet was re-suspended in Nuclei Extraction Buffer (0.5% NP40,0.25% Triton X-100, 10mM Tris-Cl pH7.5, 3mM CaCl2, 0.25mM Sucrose, 1mM DTT, phosphatase cocktail inhibitor 2, phosphatase cocktail inhibitor 3). A 3% *Drosophila* S2 cell spike-in was added at this point. This was followed by 1% formaldehyde fixation for 7mins, which was quenched with the addition of glycine to a final 125mM concentration. The nuclei pellet was re-suspended in SDS lysis buffer (1% SDS, 50mM Tris-Cl pH8.0, 10mM EDTA) and incubated on ice, followed by the addition of ChIP dilution buffer. Samples were sonicated and 1/10^th^ volume of Triton X-100 was added to dilute SDS in the solution. Samples were pellet, and supernatant was collected that contained the sonicated chromatin. Test de-crosslinking was performed on 1/8^th^ of the sonicated chromatin, and analysed using a TapeStation DNA HS tape to verify the DNA fragment range was between 100-500bp. Commercial *Drosophila* S2 chromatin (Active Motif 53083) spike-in was added at this point (at 3% of the amount of amount of *S. mediterranea* prepared chromatin) if S2 cells had been added earlier before chromatin preparation.

Protein A-covered Dynabeads were used for immunoprecipitation (IP). 50μl of Dynabeads were incubated overnight at 4c with 7μg of antibody (H3K4me3 Abcam ab8580; H3K36me3 Abcam ab9050; H3K4me1 Abcam ab8895; H3K27me3 Abcam ab6002; RNAPII-Ser5P ab5131) diluted in 0.5% BSA/PBS. Following incubation, Dynabeads were washed with 0.5% BSA/PBS, and ¼ of the total isolated chromatin was added per IP. Following overnight incubation, washes were performed 6 times with RIPA buffer (50mM HEPES-KOH pH 8.0, 500mM LiCl, 1mM EDTA, 1% NP-40, 0.7% DOC, protease inhibitors). Dynabeads were washed with TE buffer and re-suspended in Elution Buffer (50mM Tris-Cl pH 8.0, 10mM EDTA, 1% SDS). Protein was separated from Dynabeads by incubating for 15mins at 65c on a shaking heating block at 1400rpm. Eluates were de-crosslinked at 65c overnight. Input chromatin (1/8^th^ of the total chromatin amount) was also de-crosslinked at this point. Following incubation, RNAseA (0.2μg) and Proteinase K (0.2μg) was added to each sample and incubated for 1hr. DNA was purified with phenol:chloroform extraction followed by ethanol precipitation. DNA is re-suspended in TE and quantified with Qubit dsDNA HS kit. NEB Ultra II kit was used for library preparation, and clean-up was performed with Agencourt Ampure XP beads. Samples were paired-end sequenced at a length of 75 nucleotides on the Illumina NextSeq.

### ChIP-seq analysis

Reads were trimmed with Trimmomatic (Bolger et al. 2014) and aligned to a concatenated file containing both our annotated *Schmidtea mediterranea* genome as well as the *Drosophila melanogaster* release 6 reference genome (Hoskins et al. 2015) using BWA mem 0.7.12 (Li and Durbin 2009). Only uniquely mapping reads were considered further. Paired reads that map to both species were also removed. Picard tools-1.115 was used to remove duplicate reads. Reads were separated into sets that mapped to *Drosophila* or *S. mediterranea* using custom python scripts (documented in IPython notebook in **Supplementary File 6**). The number of reads aligning to the *Drosophila* genome were calculated for use in normalization calculations. For each paired or single map read, coordinates representing 100bp at the centre of sequenced were parsed and written to a BED file.

The genomecov function was used in BEDTools 2.27.0 (Quinlan and Hall 2010) to generate coverage tracks in bedgraph format. The resultant bedgraph file was converted to bigwig format UCSC’s bedgraphtoBigWig tool (Kent et al. 2010). Deeptools2’s computeMatrix was used to extract coverage around 2.5kb or 5kb either side of the annotated TSS for each annotated locus in 50bp windows for each sample and corresponding input (Ramírez et al. 2016). A normalization factor was calculated using the number of mapped reads corresponding to the *Drosophila* spike-in to control for between IP technical variation (Orlando et al. 2014). A scaling factor for input ChIP-seq libraries was calculated using the deepTools2 python API that uses the SES method (Diaz et al. 2012). The mean normalized coverage was calculated for each sample and input. The normalized input coverage was subtracted from the normalized sample coverage to generate a final coverage track for downstream visualization and analyses. Individual gene profiles for given ChIP-seq tracks could then be visualised and sequences for those genes plotted in this paper are given in **Supplementary File 7**.

To calculate the correlation of ChIP-seq signal coverage to proportional FACS expression, two vector values were calculated. The first vector was proportional FACS expression for all genomic loci. The second vector was ChIP-seq coverage at each 50bp position 2.5kb either side of the TSS. A Spearman’s rank correlation was performed on both vectors yielding a correlation value for the assayed position. The correlation value for each non-overlapping 50bp window was then plotted on a graph.

For comparison of profiles between different epigenetic marks a percentage coverage was calculated for each mark. The maximum coverage was found across all 5kb or 10kb regions for all loci. Absolute normalized coverage for each 50bp window was then divided by the maximum coverage observed for that mark in the genome, resulting in a percentage coverage in each 50bp window for each mark.

For calculation of pausing index, we added normalized coverage to input 500bp either side of the annotated TSS for each gene and divided this value by the total coverage between 500bp and 2.5kb downstream of the TSS.

Detailed methods for ChIP-seq analysis are documented in **Supplementary File 6**.

#### Data access

ChIP short read data have been deposited in the NCBI BioProject under the accession (BioProject; https://www.ncbi.nlm.nih.gov/bioproject/) numbers PRJNA471851 and PRJNA338116. Annotations made on the *S. mediterranea* genome and used in this study are available as compressed GFF file (**Supplementary File 8**)

## Acknowledgements

This work was funded grants from th the MRC (MR/M000133/1) and BBSRC (BB/K007564/1) awarded to AA. AD is funded by a BBSRC studentship (BB/J014427/1).

## Author contributions

AAA originally conceived and designed the study, upon which AD innovated. AD, YM, and PA performed ChIP-seq experiments. SH, SS, and AL assisted with optimization of ChIP-seq and RNA-seq protocols. AD and DK performed bioinformatic analyses. AD, DK and AAA wrote, reviewed, and revised the manuscript.

